# The extended recovery ring stage survival assay provides superior prediction of patient clearance half life and increases throughput

**DOI:** 10.1101/846329

**Authors:** Sage Z. Davis, Puspendra P. Singh, Katelyn M. Vendrely, Douglas A. Shoue, Lisa A. Checkley, Marina McDew-White, Katrina A. Button-Simons, Zione Cassady, Mackenzie A.C. Sievert, Gabriel J. Foster, François H. Nosten, Timothy J.C. Anderson, Michael T. Ferdig

## Abstract

**Background:** Tracking and understanding artemisinin resistance is key for preventing global setbacks in malaria eradication efforts. The ring-stage survival assay (RSA) is the current gold standard for *in vitro* artemisinin resistance phenotyping. However, the RSA has several drawbacks: it is relatively low throughput, has high variance due to microscopy readout, and correlates poorly with the current benchmark for *in vivo* resistance, patient clearance half-life post-artemisinin treatment. Here a modified RSA is presented, the extended Recovery Ring-stage Survival Assay (eRRSA), using 15 cloned patient isolates from Southeast Asia with a range of patient clearance half-lives, including parasite isolates with and without *kelch13* mutations.

**Methods:** *P. falciparum* cultures were synchronized with single layer Percoll during the schizont stage of the erythrocytic cycle. Cultures were left to reinvade to early ring-stage and parasitemia was quantified using flow cytometry. Cultures were diluted to 2% hematocrit and 0.5% parasitemia in a 96-well plate to start the assay, allowing for increased throughput and decreased variability between biological replicates. Parasites were treated with 700nM of dihydroartemisinin or an equivalent amount of dimethyl sulfoxide (DMSO) for 6 h, washed three times in drug-free media, and incubated for 66 or 114 h, when samples were collected and frozen for PCR amplification. A SYBR Green-based quantitative PCR method was used to quantify the fold-change between treated and untreated samples.

**Results:** 15 cloned patient isolates from Southeast Asia with a range of patient clearance half-lives were assayed using the eRRSA. Due to the large number of pyknotic and dying parasites at 66 h post-exposure (72 h sample), parasites were grown for an additional cell cycle (114 h post-exposure, 120 h sample), which drastically improved correlation with patient clearance half-life compared to the 66 h post-exposure sample. A Spearman correlation of 0.8393 between fold change and patient clearance half-life was identified in these 15 isolates from Southeast Asia, which is the strongest correlation reported to date.

**Conclusions:** eRRSA drastically increases the efficiency and accuracy of *in vitro* artemisinin resistance phenotyping compared to the traditional RSA, which paves the way for extensive *in vitro* phenotyping of hundreds of artemisinin resistant parasites.

## Background

Artemisinin (ART) resistance in malaria parasites is rapidly spreading through Southeast Asia, and recent reports indicate that resistance has reached Southern Asia (1–3). As artemisinin combination therapies (ACTs) are the recommended course of treatment for uncomplicated malaria by the World Health Organization (WHO), the rapid rise and spread of ART resistance raises concerns for the future of malaria treatment (4). The ability to track and understand ART resistance will be key in preventing global setbacks in malaria eradication efforts.

Measuring ART resistance is typically done *in vivo* using patient clearance half-life (PC_1/2_), an assay that measures the linear decline of parasitemia in patients after drug treatment (5–7). Clinical ART resistance manifests as a delayed clearance of parasites from a patient’s blood following treatment and is defined as a PC_1/2_ ≥ 5 h (7). While the PC_1/2_ provides a method to track ART resistance in the field, it has drawbacks, namely the requirement for patients to meet a strict inclusion criteria and agree to hospitalization to measure the PC_1/2_ (8). To avoid this costly measure*, in vitro* measures of ART resistance have been developed. One of the most common *in vitro* measures of antimalarial drug resistance is the 50% inhibitory concentration (IC_50_), which exposes parasites to serial dilutions of drug. However, delayed parasite clearance (as measured by PC_1/2_) is not associated with a significant change in ART IC_50_ (8–10). This is because later parasite stages (such as trophozoites and schizonts) are highly susceptible to ART, but early ring-stage ART resistant parasites (0-3 h) are able to survive pulses of ART. Therefore, the ring-stage survival assay (RSA) was developed to distinguish ART resistant parasites *in vitro* and to have a better correlation with PC_1/2_ data than artemisinin IC_50_s (6, 8, 11).

The RSA has been the gold standard for measuring ART resistance *in vitro*, but it is a multi-step, laborious, and time-consuming assay that requires high volumes of very synchronized parasites. It is essential that the parasites are tightly synchronized in order to assay during the short window (0-3 h) that can differentiate ART resistant parasites from ART sensitive parasites. To do this, both a Percoll gradient and sorbitol are typically used (8), but several alterations have been attempted to increase the throughput of the assay such as using both sorbitol and magnet columns (12), using syringe filters to select for merozoites (13), and using a dual layer Percoll gradient, as has been done previously in other malaria assays (14, 15). Another major bottleneck and source of variability in the final readout of the RSA is counting viable malaria parasites by microscopy (8, 11, 14). To increase throughput, flow cytometry has become heavily utilized as an alternative to counting viable parasites by microscopy, removing hours of counting slides and human error (14, 16). However, staining of cells for flow cytometry to detect viable parasites is time sensitive and requires samples to be prepared immediately after the 66 h incubation, which can be time consuming and inconvenient (14).

Despite these advances in the protocol, the RSA is still far from being both high-throughput and highly reflective of PC_1/2_. Recently, Mukherjee *et al.* used the RSA to measure the percent survival of 36 culture-adapted parasites, but only showed a correlation with PC_1/2_ data of 0.377, suggesting there is still significant room for improvement (Spearman’s Rho, internal calculations based off of supplemental data) (17).

Here a modified RSA is presented: the extended ring-stage recovery assay (eRRSA). This modified RSA protocol utilizes a simple single layer Percoll synchronization, flow cytometry to determine the stage and parasitemia for assay setup, a 96-well plate format for the assay, and a SYBR Green-based quantitative PCR (qPCR) method as the final readout. These modifications allow for a high-throughput *in vitro* experiment that better reproduces PC_1/2_, allowing for improved segregation of resistant and sensitive parasites, as well as improved sorting of moderately resistant parasites. Further, efficiency improvements in the eRRSA allow for high throughput *in vitro* testing of ART resistance, accelerating our understanding of artemisinin resistance in the lab and providing a more accurate method to track the spread of resistance.

## Methods

### Parasite isolates

To evaluate the eRRSA methods, *Plasmodium falciparum* isolates with varying *kelch13* mutations and PC_1/2_ were chosen. These isolates were derived from cloning by limiting dilution from patient samples. A total of 15 parasite isolates were chosen, 9 of which have *kelch13* mutations (including one C580Y mutant, the most common *kelch13* mutation found in Southeast Asia currently), and a PC_1/2_ distribution between 1.67 and 9.24. All 15 parasite isolates were isolated from patients on the Thailand-Myanmar border between 2008 and 2012. 3D7 was used as a control for comparison to the 15 Southeast Asian isolates (Table 1) (18, 19).

**Table 1:**
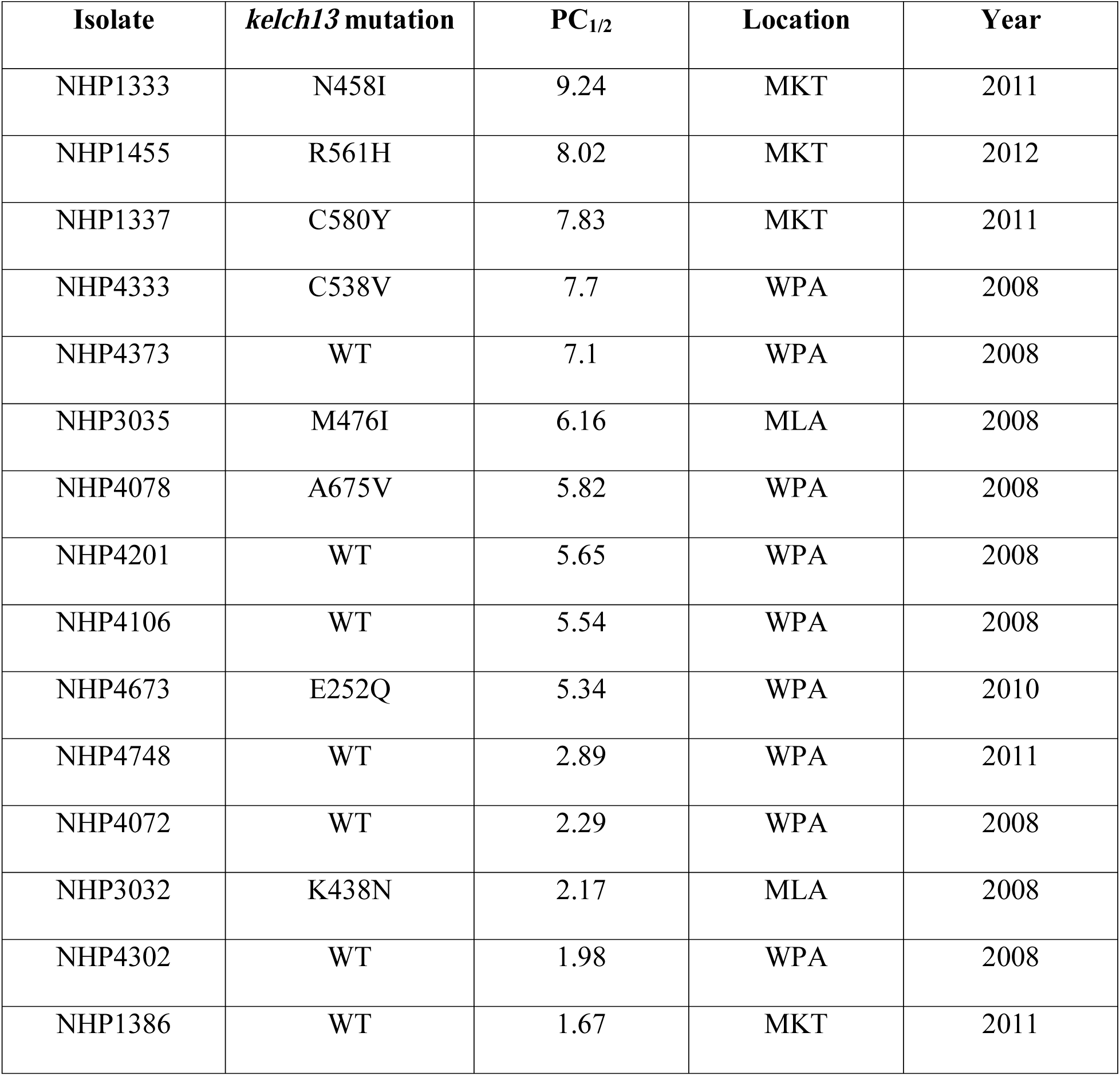
Parasite isolates used in this study. Overview of 15 parasite isolates used in this study. Patient clearance rates were calculated with (5). Location of isolate collection and year of collection are as reported from Phyo et al. (19), Taylor et al. (24), and Cheeseman et al. (25). Samples were all collected from clinics on the northwestern border of Thailand. MKT = Mawker Thai, MLA = Maela, and WPA = Wang Pha.

### Parasite culture

*P. falciparum* isolates were cultured using standard methods in human red blood cells (RBC) (Biochemed Services, Winchester, VA and Interstate Blood Bank, Memphis, TN) suspended in complete medium (CM) containing RPMI 1640 with L-glutamine (Gibco, Life Technologies.), 50 mg/L hypoxanthine (Calbiochem,Sigma-Aldrich), 25 mM HEPES (Corning, VWR, 0.5% Albumax II (Gibco, Life Technologies.), 10 mg/L gentamicin (Gibco, Life Technologies) and 0.225% NaHCO_3_ (Corning, VWR) at 5% hematocrit. Cultures were grown separately in sealed flasks at 37°C under an atmosphere of 5% CO_2_/5% O_2_/90% N_2_.

### Percoll synchronization

Parasites were synchronized using a density gradient method as previously described with slight modifications (14, 15, 20). Briefly, 350ul of packed, infected erythrocytes at high schizogony (>50% schizonts) was suspended in 2ml of RPMI. Cultures were layered over a single 70% Percoll (Sigma-Aldrich) layer in 1x RPMI and 13.3% sorbitol in phosphate buffer saline (PBS) and centrifuged (1561x*g* for 10 min, no brake). The top layer of infected late stage schizonts was then removed and washed with 10 ml of RPMI twice. Cultures were then suspended in 2 ml of CM at 2% hematocrit and placed in culture flasks on a shaker in a 37 °C incubator for 4 h to allow for re-invasion.

### Flow cytometry

4 h after Percoll synchronization (unless noted otherwise), samples were measured by flow cytometry as previously described with slight modifications to determine parasitemia (14, 16). Briefly, 80 μl of culture and an RBC control incubated for at least 8h at 5% hematocrit in CM were stained with SYBR Green I (SYBR) and SYTO 61 (SYTO) and measured on a guava easyCyte HT (Luminex Co.). Analysis was performed with guavaSoft version 3.3 (Luminex Co.). 50,000 events were recorded for both the RBC control and samples to determine relative parasitemias.

### eRRSA setup

2 h post-cytometric quantitation (or 6 h after Percoll synchronization) samples whose stage composition was >70% rings as determined by flow cytometry were diluted to 2% hematocrit and 0.5% parasitemia (unless otherwise noted), and 200 μl of culture was aliquoted into 6 wells of a flat bottom 96-well plate. Each treated and untreated sample had three technical replicates: RBC controls were aliquoted into 2 wells at 2% hematocrit and 200 μl. 3 wells of parasites and 1 well of the RBC control were treated with 700nM dihydroartemisinin (DHA) (Sigma-Aldrich); an additional 3 wells of parasites and 1 well of RBC control was treated with an equivalent amount of dimethyl sulfoxide (DMSO) (ThermoFisher) as untreated controls. Parasites were incubated for 6 h, and then washed three times with 150 μl of RPMI to remove drug. Samples were then suspended in CM and placed back in the incubator. 66 h after drug removal, 20 μl of sample from each well was collected and frozen for qPCR amplification (72 h sample). Plates were then placed back in the incubator for another 48 h, after which 20 μl of sample was again collected and frozen for qPCR amplification (120 h sample).

### qPCR Amplification

qPCR amplification was done as previously reported, with slight modifications (21). Ring stage samples were quantified at 72 and 120 h post-drug treatment. qPCR was performed using the Phusion Blood Direct PCR kit (ThermoFisher, cat # F547L), supplemented with 1x SYBR. 3 μl of culture was used in a 10 μl reaction and amplified using forward and reverse primers of the *Pfcrt* gene. PCR amplification was measured using the fast mode of the ABI 7900HT, with a 20 s denaturation at 95 °C, followed by 30 cycles of 95 °C for 1 s, 62.3 °C for 30 s, and 65 °C for 15 s (Additional file 1). Cycle threshold (Ct) values were calculated using the ABI SDS 2.4.1. Fold change (2^ΔCt^) was calculated by determining the average ΔCt for the three technical replicates for the untreated and treated samples by applying the following equation:

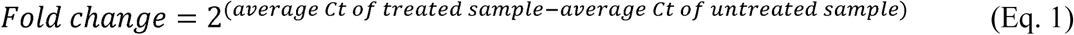

All statistics were performed and figures generated using GraphPad Prism version 8.2.1.

## Results

### Using a SYBR Green-based quantitative PCR method to quantify the fold-change between treated and untreated samples

Percent proliferation is the standard RSA measurement to determine whether a parasite is artemisinin resistant or sensitive. This is calculated by dividing the percent parasitemia in the treated (DHA) sample over the percent parasitemia in the untreated (DMSO) sample. Parasitemia is determined either by counting the number of viable and nonviable parasites using blood smears and microscopy or by flow cytometry. Determining parasitemia with microscopy is cost effective and convenient but is also highly variable and time-consuming. Flow cytometry is typically very accurate; however, the process must be begun at upon reaching a timepoint, adding a substantial time investment at the point of sampling. To find another measurement of parasitemia that could be automated and have less variability, qPCR on parasite genomic DNA was tested. A standard curve of percent parasitemia (as measured by flow cytometry) shows excellent inverse correlation with Ct values as measured by qPCR (Figure 1). To quantify the difference between treated and untreated final RSA samples, fold change (2^ΔCt^) was calculated according to equation 1.

**Fig. 1.**
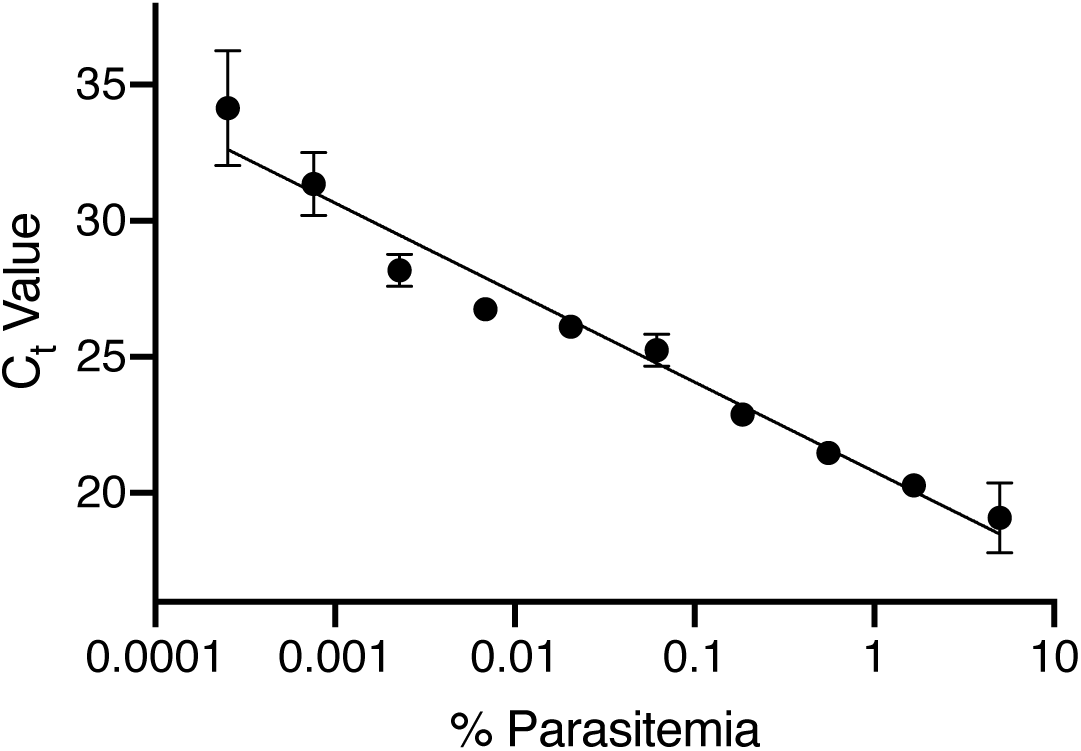
qPCR standard curve for detecting parasitemia of *P. falciparum* infected RBCs. A standard curve of samples ranging from 0.0001% parasitemia to 9% parasitemia were measured using qPCR amplification on the ABI 7900HT. Ct values were calculated based on three technical replicates using the ABI SDS 2.4.1. Ct value is inversely related to percent parasitemia (R^2^ = 0.9699) and therefore can be used as a measurement of percent parasitemia in final RSA samples.

### RSA readout at 120 h provides superior differentiation between sensitive and resistant isolates

The standard RSA determines parasite viability at 72 h after drug treatment. However, a common problem in the final readout (using either microscopy, flow cytometry, or qPCR) is the difficulty in differentiating between pyknotic (nonviable) parasites and viable parasites 72 h after drug treatment. It is also difficult to measure viable parasitemia when it can be as low as 0.01% (or even 0% in some cases), especially when measuring ART sensitive parasites (22). To address these issues, the time to readout was extended by an additional intraerythrocytic development cycle (48 h). This extension was added to both allow parasites an additional expansion cycle, creating larger differences to distinguish resistant and sensitive isolates, and to allow erythrocytes to clear pyknotic parasites. This additional cycle provides a much greater separation between resistance and sensitive parasite isolates (Figure 2).

**Fig. 2.**
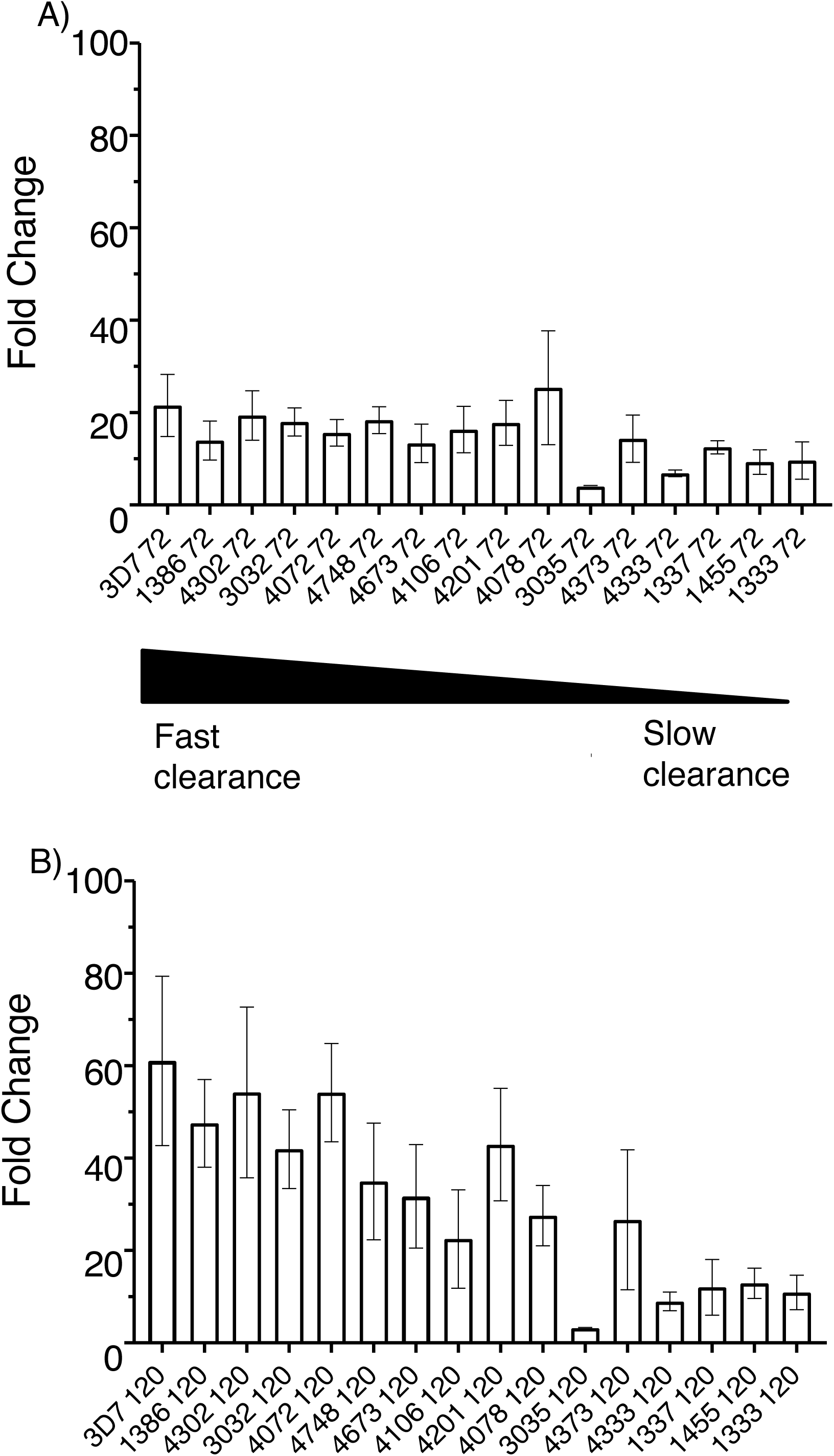
Comparison of 72 h and 120 h perturbations. Parasites were set-up using the eRRSA protocol: 0.5% parasitemia at early ring stage and 700 nM DHA was applied and washed off after 6 h. Samples were collected at (A) 72 h post-drug treatment and at (B) 120 h post-drug treatment. Parasites are ordered from the smallest PC_1/2_ (least resistant) to the largest PC_1/2_ (most resistant) from left to right (with the sensitive control, 3D7 on the far left). Sensitive and resistant parasites were more distinguishable based on their 120 h post-drug treatment sample fold changes.

### Assay set-up conditions have a substantial impact on RSA outcomes

The RSA is a growth assay targeted at a very narrow window of the parasite intraerythrocytic development cycle. In order to maximize the precision of the assay, the growth and the timing of the target window were carefully optimized. First, as it has been established that growth rates can vary based on parasitemia, the effect of varying starting parasitemias on RSA outcome was observed (23). RSA was performed on three parasite isolates (3D7, 1337, and 4673) at varying starting parasitemias as determined by a microscopist, and samples were collected at 72h and 120h (Additional file 2A and 2B, respectively). For the three parasite isolates tested, 3D7 was the ART sensitive control and 1337 and 4673 were two ART resistant parasite isolates as determined by their PC_1/2_ values (1337 PC_1/2_ = 7.84 and 4673 PC_1/2_ = 5.34). The 0.25% starting parasitemia showed the most distinguishable phenotype between the ART sensitive 3D7 control and the two ART resistant isolates at 120 h (Additional file 2B). It was noted internally that determinations of parasitemia by microscopy varied widely and underestimated parasitemia compared to flow cytometry, likely due to the difficulty of correctly identifying new invasions. A comparison of RSA results from isolates set up at 0.25% parasitemia determined by microscopy and 0.5% parasitemia determined by flow cytometry showed no difference between the two (Additional file 3). As a result, in subsequent set-ups starting parasitemia was determined by flow cytometry and was normalized to 0.5% parasitemia.

A key factor in the RSA is applying the drug treatment in the tight 0-3 h window of the parasite life cycle that can differentiate ART resistant parasites from ART sensitive parasites. Therefore, the time from Percoll synchronization to drug treatment was also varied to find the optimal time for drug treatment. The same three parasite isolates, (3D7, 1337, and 4673), were set up and treated with 700nM DHA 4 h, 6h, 8 h, or 10 h after Percoll synchronization and samples collected at 72 h and 120 h post-drug treatment (Additional file 2C and 2D, respectively). All parasites were set up at a starting parasitemia of 0.5% as measured by flow cytometry. The time that resulted in the most consistent and distinguishable phenotypes between the ART sensitive and resistant isolates was 6 h post-Percoll synchronization. Controlling these factors resulted in a more consistent and reproducible RSA phenotype.

### Defining the eRRSA as an improvement over the standard RSA

Based on the previously described optimizations, the eRRSA is defined as a single layer Percoll synchronization, flow cytometry measurement of parasite parasitemia and stage prior to assay set-up, 700nM DHA treatment of 200ul of parasites at 2% hematocrit and 0.5% parasitemia in 96-well plates that are set up at 6 h post-Percoll synchronization, drug washed off 6 h after application, and samples for qPCR readout collected at 120 h post-drug treatment (Additional file 4). This protocol provides the most consistent and high-throughput results. These modifications made to the standard RSA are a new *in vitro* ART resistance phenotyping method: the Extended Ring-stage Recovery assay (eRRSA).

### eRRSA correlates better with PC_1/2_ than RSA

The RSA was introduced as an *in vitro* method that better captures the gold standard for *in vivo* ART resistance, PC_1/2_, than the traditional IC_50_. For a new assay to be relevant, it should perform at least comparably to the existing standard. Therefore, the eRRSA was used to assay 15 isolates from Southeast Asia with varying known PC_1/2_ and *kelch13* mutations collected between 2008 and 2012 (Table 1). We collected at least three biological replicates (each with three technical replicates) for each isolate and collected samples at 72 h and 120 h post-drug treatment and compared the viability of treated and untreated parasites at each stage.

The fold change data was then compared to PC_1/2_: at the 72 h timepoint across 15 isolates, the eRRSA has a Spearman correlation coefficient of −0.6071 (Figure 3A). This is comparable to other RSA correlations in the field, showing that the improvements made to the RSA protocol do not significantly affect the outcome while increasing efficiency and ease of the assay (8, 17). The 120 h correlations, however, improved Spearman correlation between fold change and PC_1/2_ to - 0.8393 (Figure 3B).

**Fig. 3.**
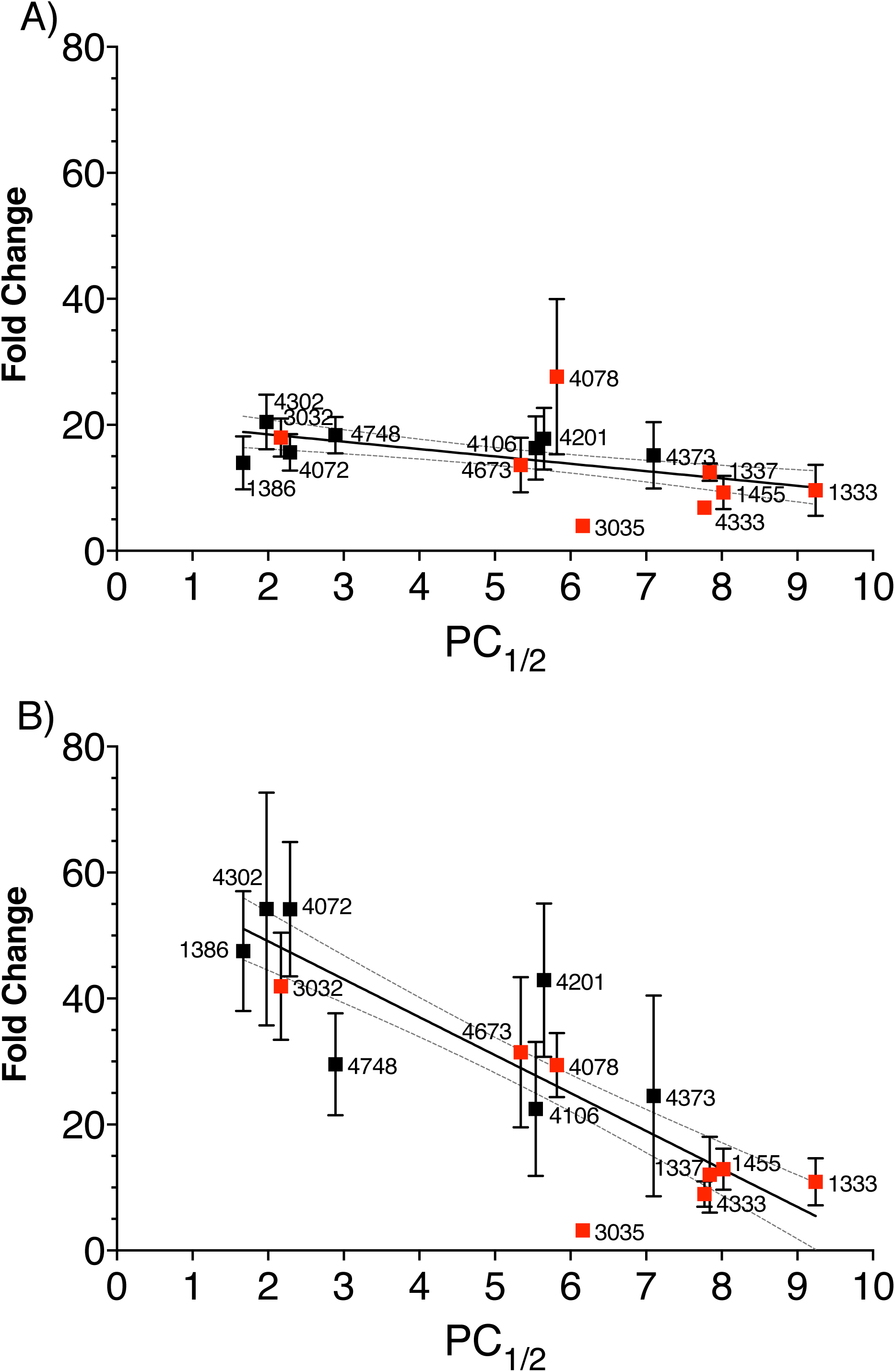
120 h eRRSA increases correlation with PC_1/2_ compared to 72 h. (A) 15 isolates from Southeast Asia with varying PC_1/2_ were assayed using the eRRSA. 72 h post-drug treatment samples were measured to give a fold change for each isolate and those fold changes were correlated with each isolate’s PC_1/2_ (Spearman r =-0.6071). Isolates with red boxes are *kelch13* mutants and the 95% confidence interval around the best-fit line is denoted with dotted lines. (B) The same 15 isolates were assayed using the eRRSA and 120 h post-drug treatment samples were measured to calculate a fold change for each isolate. The fold changes were correlated with PC_1/2_ (isolates marked with red boxes are *kelch13* mutants and the 95% confidence interval of the best-fit line is denoted with dotted lines) (Spearman r = −0.8393); the 120 h eRRSA samples increased correlation with PC_1/2_ compared to the 72 h samples.

## Discussion

With the gradual spreading of artemisinin resistance throughout Southeast Asia, it is imperative that resistance can be accurately measured both in the field and in the lab. To date, the RSA has been the golden standard for *in vitro* measurement of artemisinin resistance. Despite this, there is a need for a more accurate, higher throughput *in vitro* measurement alternative to the current RSA to accelerate our understanding of artemisinin resistance. We have developed the eRRSA for this purpose and have shown that it can outperform the RSA in both accuracy and throughput.

A major bottleneck and source of variability in the RSA is the final readout to determine the ratio of viable parasites in the treated and untreated cultures. RSA uses microscopy or flow cytometry as the final readout, while the eRRSA uses qPCR. When using microscopy to compare viable and nonviable parasites, the presence of the ring-like structure of healthy, viable parasites are compared to the collapsed, nonviable parasites which can be highly subjective and time consuming, as shown by the original RSA paper which required two to three microscopists (14). Flow cytometry utilizes the DNA stain SYBR and the mitochondrial stain MitoTracker red to differentiate between viable and nonviable parasites. This eliminates the requirement of microscopists and drastically decreases the labor required to determine percent proliferation of parasites (14, 16). However, staining of cells for flow cytometry is time sensitive and must be done immediately following the end of the RSA (at 72 h), which can limit flexibility and lengthens an assay that already demands long hours. qPCR measure viable parasites solely by concentration of genomic material, comparing the efficiency of parasites to proliferate post-artemisinin perturbation. Here, the efficacy of qPCR is demonstrated as a readout for proliferation in a survival assay context. The use of qPCR allows for both smaller sample sizes and a delayed readout, rendering the protocol easier and more precise.

The eRRSA measures the difference in genomic content between treated and untreated malaria parasites at 120 h post perturbation, a full 48 h (or a full life cycle) after the RSA collections. Our results demonstrate that by allowing parasites an extra life cycle to recover, the differences between resistant and sensitive parasites are made even more drastic, suggesting the extra lifecycle for recovery helps further differentiate viable and nonviable parasites. With an extra life cycle for recovery from drug treatment, the eRRSA measures recovery of the parasites, rather than the survivability of the parasites after treatment, and provides a more consistent readout compared to other publications (Additional file 5).

The demonstration of the effect of various setup conditions on the final outcome required an optimization of these parameters in the eRRSA. With no sorbitol synchronization required and only one single layer Percoll synchronization to select schizonts, parasites are synchronized easier, quicker, and closer to the ring-stage so that they can be set up in the assay sooner to avoid losing their synchronization. The variability in assay outcome caused by varying the delay between synchronization and treatment is likely due to the short window (0-3 h) that differentiates ART resistant parasites from ART sensitive parasites. The eRRSA assay is set up 6 h post-Percoll, which allows for a high percentage of early ring-stage, tightly synchronized parasites. Using flow cytometry for parasitemia measurements post-Percoll synchronization allows for rapid and accurate parasitemia determination and staging for many parasites at once at the set-up of the assay, which is an essential factor for the results of both RSA and eRRSA. Starting parasitemia has a substantial effect on growth throughout the assay; by using a lower starting parasitemia (0.5%) compared to the *in vitro* RSA, the eRRSA has a lower volume and parasitemia requirement while also allowing for more precise measurement of growth with and without drug treatment. Lower culture volume requirements permit the use of 96-well plates, which uses less reagents, time, and space.

Finally, it was demonstrated that the eRRSA shows superior correlation with the clinical phenotype PC_1/2_. As ART resistance is currently presenting as a continuous phenotype, the ability to accurately determine intermediate phenotypes is critical in understanding ART resistance and identifying contributing genotypes beyond K13 propeller mutations. The efficacy and efficiency of the eRRSA makes it an excellent replacement for the traditional RSA in any study of ART resistance requiring accuracy and high-throughput.

The 15 cloned parasites examined include 3 which lack K13 mutations but show PC_1/2_ >5. These include one clone (NHP4373) with PC_1/2_ = 7.1. Interestingly, these clones also show high eRRSA values, confirming their ART-R status. These results provide further support that ART-R may result from mutations elsewhere in the parasite genome, or perhaps from non-coding regulatory changes controlling K13 activity (17).

## Conclusions

The eRRSA method described here provides a more robust *in vitro* representation of PC_1/2_ while also providing vastly improved throughput. Widespread adaptation of the eRRSA should significantly accelerate our understanding of artemisinin resistance, allowing for both high throughput surveillance of the spread of resistance and for the precise phenotyping necessary to uncover complex genetic contributors to resistance.

## List of abbreviations

RSA: Ring-stage survival assay
eRRSA: extended Recovery Ring-stage Survival Assay
DMSO: Dimethyl sulfoxide
ART: Artemisinin
ACTs: Artemisinin combination therapies
WHO: World Health Organization
PC_1/2_: Patient clearance half-life
qPCR: quantitative Polymerase Chain Reaction
RBC: Red blood cell
CM: Complete media
PBS: Phosphate buffer saline
SYBR: SYBR Green I
SYTO: SYTO 61 red fluorescent nucleic acid stain DHA Dihydroartemisinin
Ct: cycle threshold
QTL: Quantitative trait loci

## Declarations

### Ethics approval and consent to participate

Ethical approval for the use of human blood in this study was granted by the Institutional Review Board of the University of Notre Dame. All of the blood used for the *in vitro* culturing of parasites was obtained from healthy adult volunteers and drawn by trained personnel from Interstate Blood Bank.

### Consent for publication

Not applicable.

### Availability of data and material

Data can be made available upon request to the corresponding author.

## Authors’ contributions

SZD, LAC, DAS, and MTF conceived and designed the experiments. PPS, LAC, DAS, GF, and MACs performed the experiments, analysed the data, and created figures. SZD, KMV, LAC, GF, and MTF wrote the manuscript. FHN, MMW and TJCA contributed materials/reagents/parasites. All authors read, edited, and approved the final manuscript.

## Acknowledgements

We would like to acknowledge members of the Ferdig lab for helpful discussions.

## Additional files

**Additional File 1.pdf:**
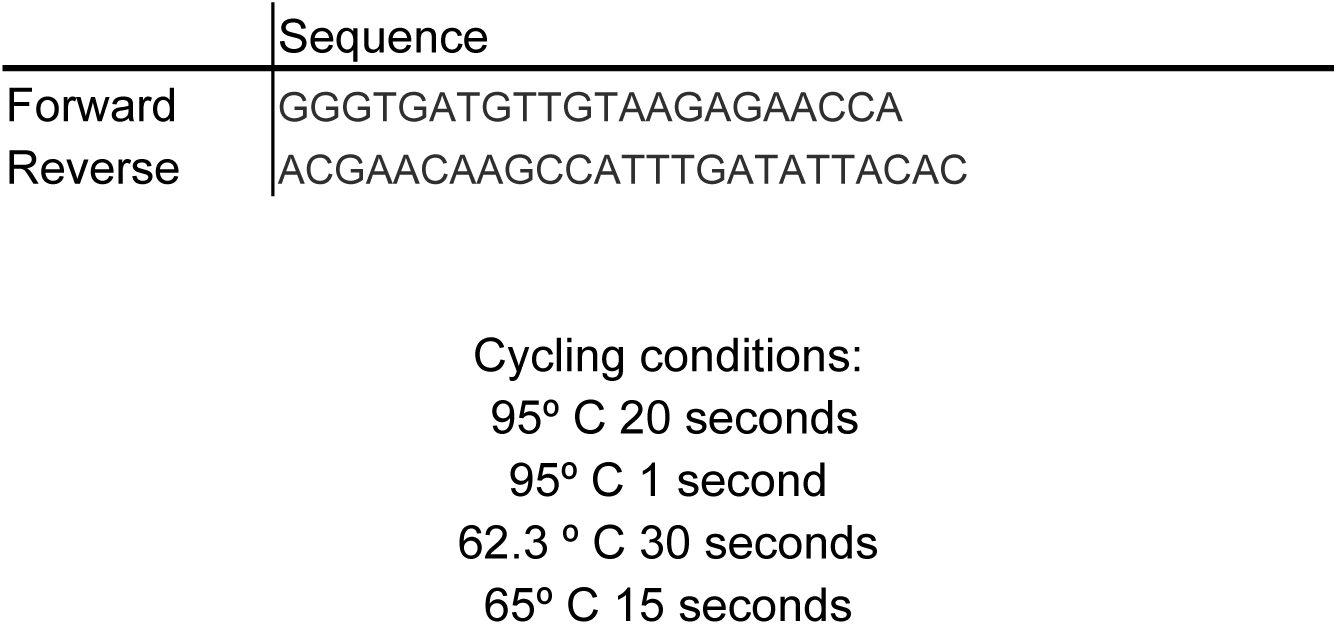
Table of primer and cycling conditions used for quantitative PCR. The forward and reverse primer sequences of the *Pfcrt* gene used in this study and the cycling conditions used in the fast mode of the ABI 7900HT.

**Additional File 2.pdf:**
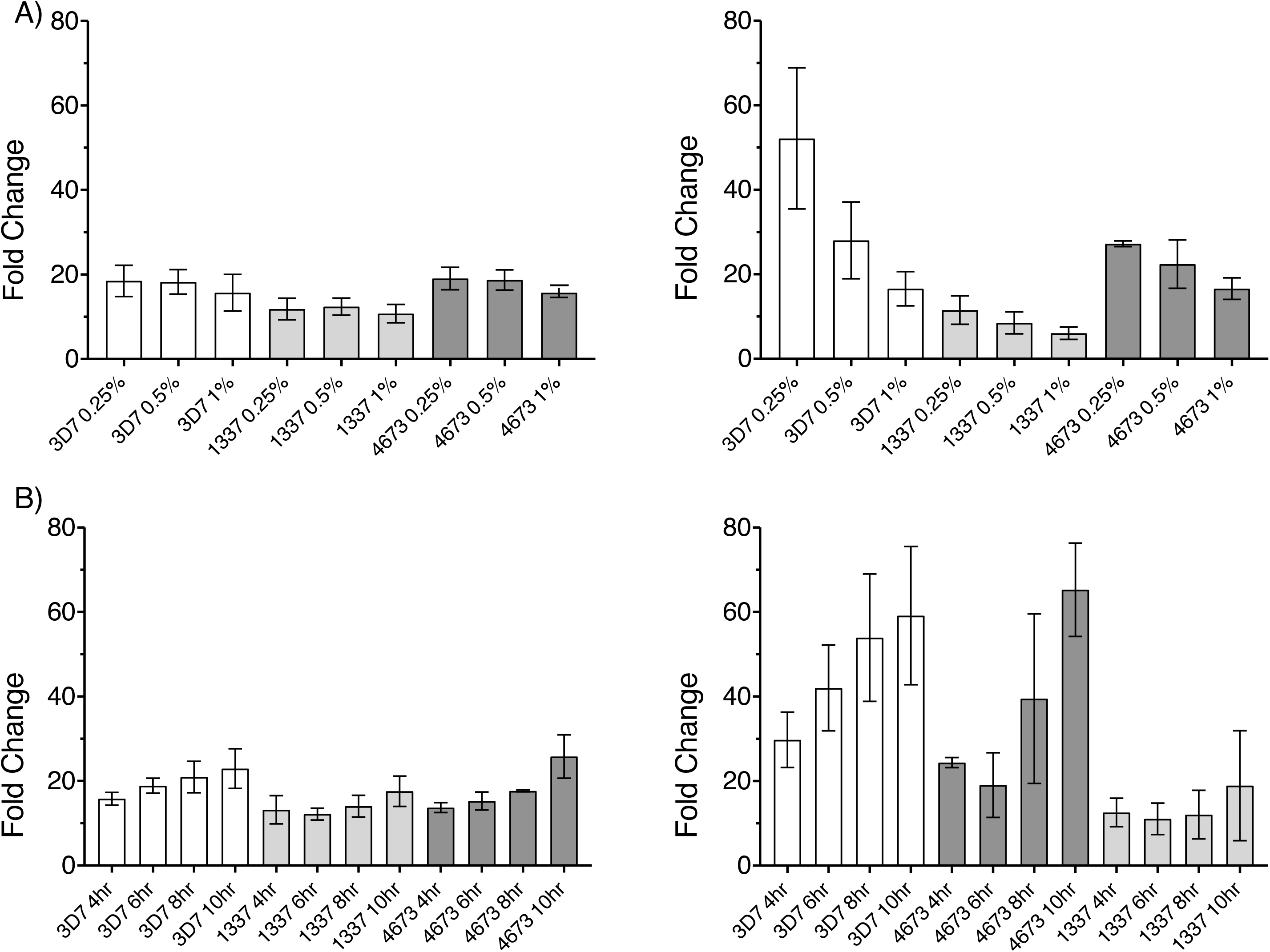
Effects of assay modification on final assay readout. (A) The starting parasitemia for the assay was set at 0.25%, 0.5%, or 1% (as measured by microscopy) for three different parasite isolates (3D7, 1337, and 4673) and the fold change between the treated (DHA) and untreated (DMSO) samples was measured with three biological replicates (each with three technical replicates) at 72 h post-drug treatment and (B) at 120 h post-drug treatment. After determining that a 0.25% parasitemia measured by microscopy was equivalent to 0.5% parasitemia measured by flow cytometry (C), the starting parasitemia was determined using flow cytometry and set to 0.5% and the time from Percoll synchronization to drug (DHA or DMSO) application was varied (4 h, 6 h 8 h, or 10 h) for the three different parasite isolates (3D7, 1337, and 4673) and the fold change between treated and untreated samples was measured with three biological replicates (each with three technical replicates) at 72 h post-drug treatment and (D) 120 h post-drug treatment.

**Additional File 3.pdf.**
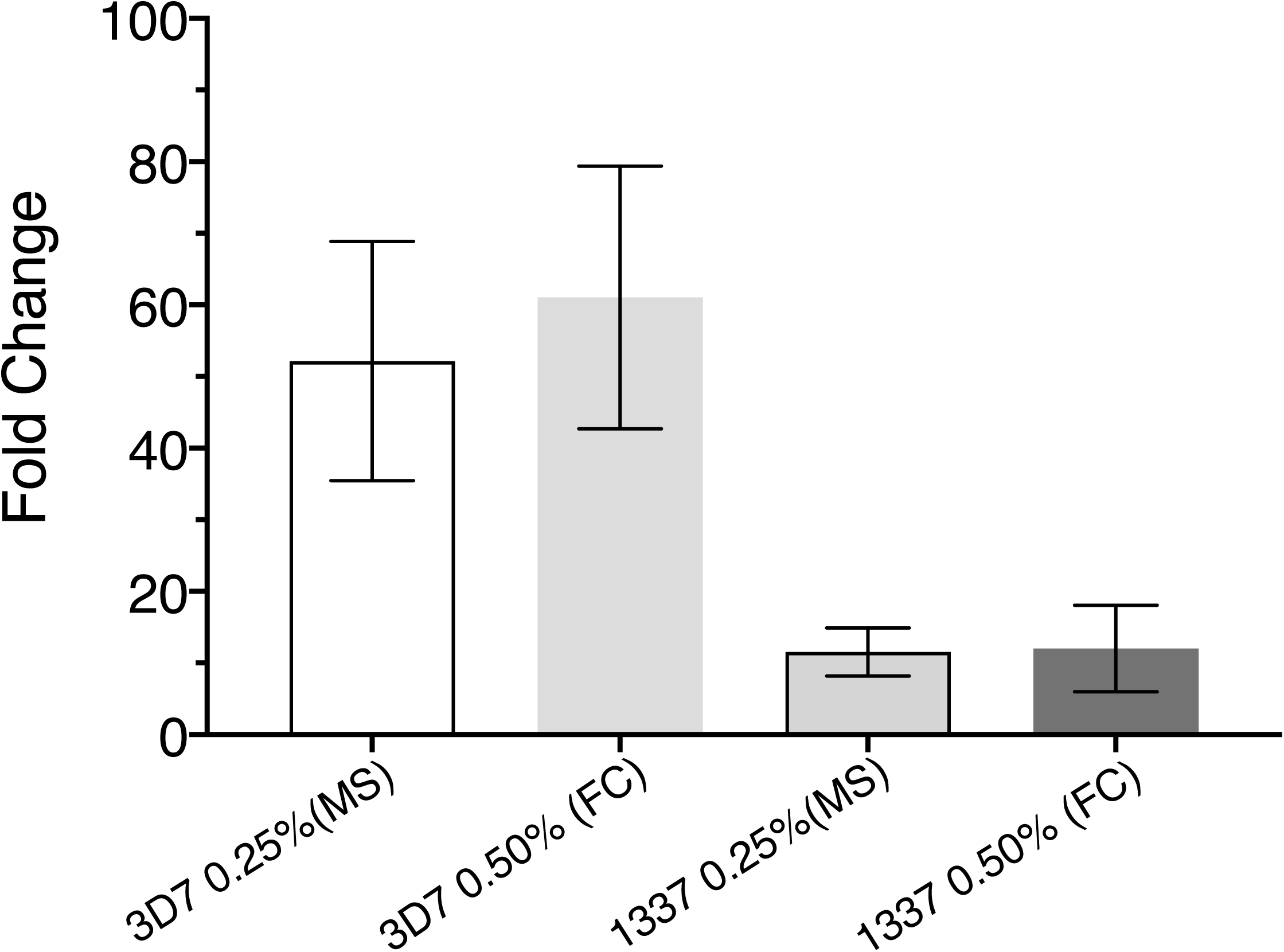
Comparison of microscopy vs flow cytometry to calculate starting parasitemia. Fold change data for two lines used as controls in eRRSA show little difference between a starting parasitemia of 0.25% by microscopy and a starting parasitemia of 0.5% determined by flow cytometry.

**Additional File 4.pdf:**
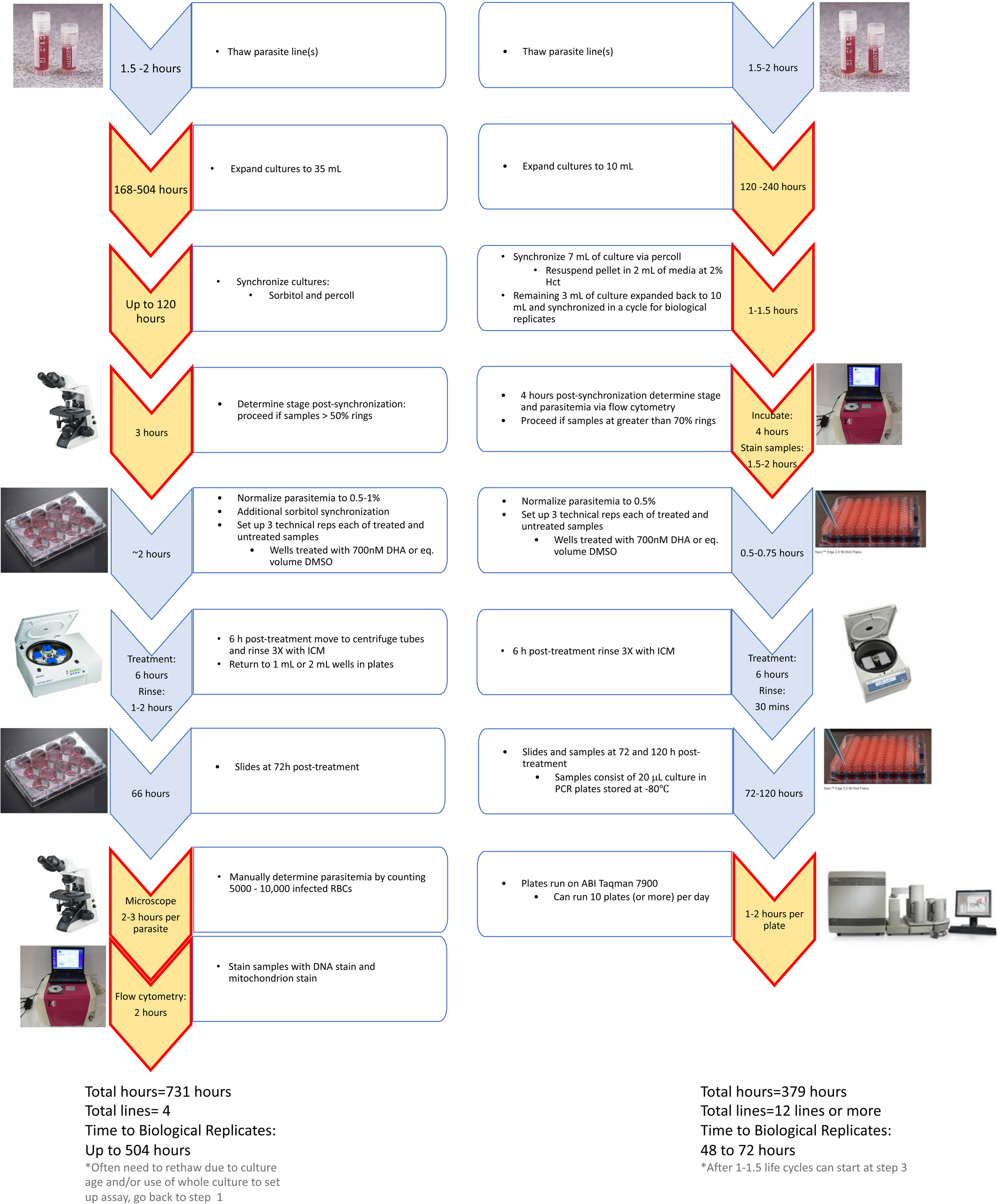
Summary of modifications for the eRRSA compared to the standard RSA. A timeline comparing the various steps and the estimated time needed for each step of the standard WWARN RSA (14) (left) and the eRRSA (right). There are limiting steps in both the eRRSA and RSA (marked with orange arrows), but the eRRSA allows for more parasite isolates with more biological replicates to be done in less time compared to the standard RSA.

**Additional File 5.pdf.**
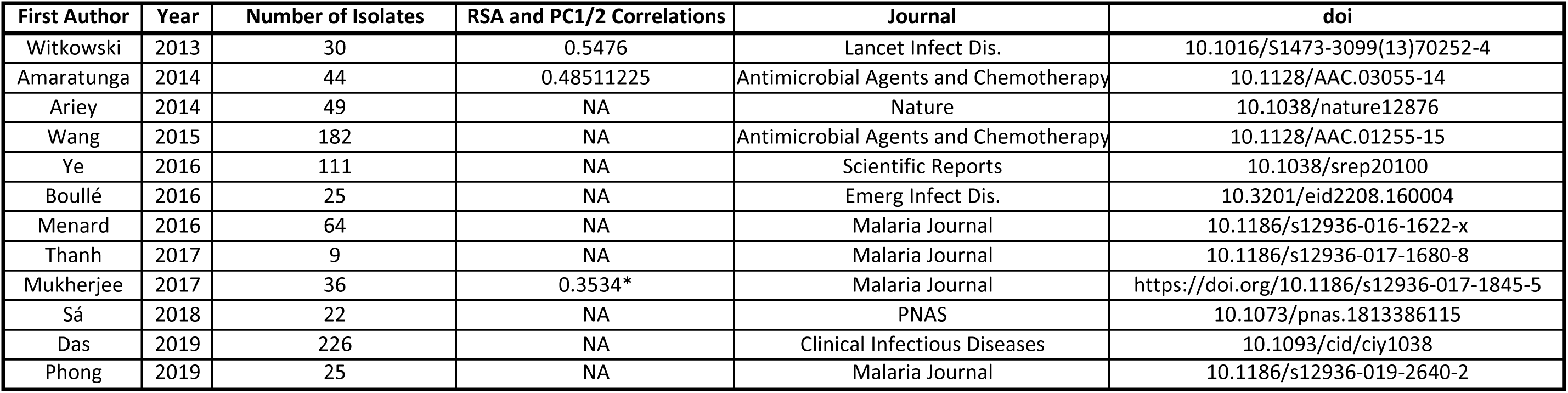
Comparison of previously published correlation rates of RSA results with PC_1/2_. Previous publications where PC_1/2_ and RSA results on the same samples were reported are listed here. If RSA results were available, then the Spearman correlation between PC_1/2_ and RSA was calculated. If the Spearman correlation was not provided, it was internally calculated and is marked with an *.

## References

1. Cerqueira GC, Cheeseman IH, Schaffner SF, Nair S, McDew-White M, Phyo AP, et al. Longitudinal genomic surveillance of Plasmodium falciparum malaria parasites reveals complex genomic architecture of emerging artemisinin resistance. Genome Biol. 2017;18(1):78.

2. Imwong M, Suwannasin K, Kunasol C, Sutawong K, Mayxay M, Rekol H, et al. The spread of artemisinin-resistant Plasmodium falciparum in the Greater Mekong subregion: a molecular epidemiology observational study. Lancet Infect Dis. 2017;17(5):491–7.

3. Das S, Saha B, Hati AK, Roy S. Evidence of Artemisinin-Resistant Plasmodium falciparum Malaria in Eastern India. N Engl J Med. 2018;379(20):1962–4.

4. Organization WH. World Malaria Report. Geneva, Switzerland: World Health Organization; 2018.

5. Flegg JA, Guerin PJ, White NJ, Stepniewska K. Standardizing the measurement of parasite clearance in falciparum malaria: the parasite clearance estimator. Malar J. 2011;10:339.

6. Group WKG-PS. Association of mutations in the Plasmodium falciparum Kelch13 gene (Pf3D7_1343700) with parasite clearance rates after artemisinin-based treatments-a WWARN individual patient data meta-analysis. BMC Med. 2019;17(1):1.

7. Ashley EA, Dhorda M, Fairhurst RM, Amaratunga C, Lim P, Suon S, et al. Spread of artemisinin resistance in Plasmodium falciparum malaria. N Engl J Med. 2014;371(5):411–23.

8. Witkowski B, Amaratunga C, Khim N, Sreng S, Chim P, Kim S, et al. Novel phenotypic assays for the detection of artemisinin-resistant Plasmodium falciparum malaria in Cambodia: invitro and ex-vivo drug-response studies. Lancet Infect Dis. 2013;13(12):1043–9.

9. Dondorp AM, Nosten F, Yi P, Das D, Phyo AP, Tarning J, et al. Artemisinin resistance in Plasmodium falciparum malaria. N Engl J Med. 2009;361(5):455–67.

10. Amaratunga C, Sreng S, Suon S, Phelps ES, Stepniewska K, Lim P, et al. Artemisinin-resistant Plasmodium falciparum in Pursat province, western Cambodia: a parasite clearance rate study. Lancet Infect Dis. 2012;12(11):851–8.

11. Klonis N, Xie SC, McCaw JM, Crespo-Ortiz MP, Zaloumis SG, Simpson JA, et al. Altered temporal response of malaria parasites determines differential sensitivity to artemisinin. Proc Natl Acad Sci U S A. 2013;110(13):5157–62.

12. Tirrell AR, Vendrely KM, Checkley LA, Davis SZ, McDew-White M, Cheeseman IH, et al. Pairwise growth competitions identify relative fitness relationships among artemisinin resistant Plasmodium falciparum field isolates. Malar J. 2019;18(1):295.

13. Kite WA, Melendez-Muniz VA, Moraes Barros RR, Wellems TE, Sá JM. Alternative methods for the Plasmodium falciparum artemisinin ring-stage survival assay with increased simplicity and parasite stage-specificity. Malaria Journal. 2016;15.

14. B Witkowski DM, C Amaratunga, RM Fairhurst. Ring-stage Survival Assays (RSA) to evaluate the in-vitro and ex-vivo susceptibility of Plasmodium falciparum to artemisinins. Institute Pasteur du Cambodge – National Institutes of Health Procedure RSAv1.2013.

15. Lewis IA, Wacker M, Olszewski KL, Cobbold SA, Baska KS, Tan A, et al. Metabolic QTL Analysis Links Chloroquine Resistance in Plasmodium falciparum to Impaired Hemoglobin Catabolism. PLoS Genetics. 2014;10:e1004085.

16. Amaratunga C, Neal AT, Fairhurst RM. Flow cytometry-based analysis of artemisinin-resistant Plasmodium falciparum in the ring-stage survival assay. Antimicrob Agents Chemother. 2014;58(8):4938–40.

17. Mukherjee A, Bopp S, Magistrado P, Wong W, Daniels R, Demas A, et al. Artemisinin resistance without pfkelch13 mutations in Plasmodium falciparum isolates from Cambodia. Malaria Journal. 2017;16.

18. Cheeseman IH, McDew-White M, Phyo AP, Sriprawat K, Nosten F, Anderson TJ. Pooled sequencing and rare variant association tests for identifying the determinants of emerging drug resistance in malaria parasites. Mol Biol Evol. 2015;32(4):1080–90.

19. Phyo AP, Nkhoma S, Stepniewska K, Ashley EA, Nair S, McGready R, et al. Emergence of artemisinin-resistant malaria on the western border of Thailand: a longitudinal study. The Lancet. 2012;379(9830):1960–6.

20. Dluzewski AR, Ling IT, Rangachari K, Bates PA, Wilson RJ. A simple method for isolating viable mature parasites of Plasmodium falciparum from cultures. Trans R Soc Trop Med Hyg. 1984;78(5):622–4.

21. Wacker MA, Turnbull LB, Walker LA, Mount MC, Ferdig MT. Quantification of multiple infections of Plasmodium falciparum in vitro. Malaria Journal. 2012;11(1):180.

22. Wang Z, Cabrera M, Yang J, Yuan L, Gupta B, Liang X, et al. Genome-wide association analysis identifies genetic loci associated with resistance to multiple antimalarials in Plasmodium falciparum from China-Myanmar border. Sci Rep. 2016;6:33891.

23. Rovira-Graells N, Aguilera-Simon S, Tinto-Font E, Cortes A. New Assays to Characterise Growth-Related Phenotypes of Plasmodium falciparum Reveal Variation in Density-Dependent Growth Inhibition between Parasite Lines. PLoS One. 2016;11(10):e0165358.

24. Taylor AR, Schaffner SF, Cerqueira GC, Nkhoma SC, Anderson TJC, Sriprawat K, et al. Quantifying connectivity between local Plasmodium falciparum malaria parasite populations using identity by descent. PLoS Genet. 2017;13(10):e1007065.

25. Anderson TJ, Nair S, McDew-White M, Cheeseman IH, Nkhoma S, Bilgic F, et al. Population Parameters Underlying an Ongoing Soft Sweep in Southeast Asian Malaria Parasites. Mol Biol Evol. 2017;34(1):131–44.

